# Identification of a regulatory pathway governing TRAF1 via an arthritis-associated non-coding variant

**DOI:** 10.1101/2022.12.15.520628

**Authors:** Qiang Wang, Marta Martínez-Bonet, Taehyeung Kim, Jeffrey A. Sparks, Kazuyoshi Ishigaki, Xiaoting Chen, Marc Sudman, Vitor Aguiar, Marcos Chiñas Hernandez, Alexandra Wactor, Brian Wauford, Miranda C. Marion, Maria Gutierrez-Arcelus, John Bowes, Stephen Eyre, Ellen Nordal, Sampath Prahalad, Marite Rygg, Vibeke Videm, Soumya Raychaudhuri, Matthew T. Weirauch, Carl D. Langefeld, Susan D. Thompson, Peter A. Nigrovic

## Abstract

TRAF1/C5 was among the first loci shown to confer risk for inflammatory arthritis in the absence of an associated coding variant, but its genetic mechanism remains undefined. Using ImmunoChip data from 3,939 juvenile idiopathic arthritis (JIA) patients and 14,412 controls, we identified 132 plausible common non-coding variants, reduced serially by SNP-seq, electrophoretic mobility shift, and luciferase studies to the single variant rs7034653 in the third intron of *TRAF1*. Genetically manipulated experimental cells and primary monocytes from genotyped donors establish that the risk G allele reduces binding of Fos-Related Antigen 2 (FRA2), resulting in reduced TRAF1 expression and enhanced TNF production. Conditioning on this variant eliminates attributable risk for rheumatoid arthritis, implicating a mechanism shared across the arthritis spectrum. These findings reveal that rs7034653, FRA2, and TRAF1 mediate a pathway through which a non-coding causal variant drives risk of inflammatory arthritis in children and adults.

## Introduction

Translating findings from genome-wide association studies (GWAS) into pathogenic understanding has proven challenging because most implicated loci do not correspond to an exonic variant ^1^. Such loci are presumed to carry one or more variants that modulate gene expression, but definitive identification of regulatory variants has proven difficult. Strategies employed have included bioinformatic prediction, fine mapping, and experimental interrogation of plausible non-coding single nucleotide polymorphisms (SNPs) through methods such as the massively parallel reporter assay and SNP-seq ^1–4^. The challenges attendant to these strategies, and to downstream experimental confirmation that a particular variant is indeed causal, are such that few non-coding variants are clearly defined. Correspondingly, mechanistic insights obtained through GWAS have often lagged behind those gained from studies of monogenic diseases ^5^.

Among the disease families closely examined by GWAS is inflammatory arthritis. The most common disease in this group, rheumatoid arthritis (RA), has been studied in tens of thousands of patients and controls, yielding over 120 loci associated with disease risk ^6–10^. One of the first loci identified using a genome-wide approach was a region on chromosome 9 containing *TRAF1* and *C5;* unlike other genes with confirmed risk associated with RA at that time (the HLA region, *PTPN22*), the TRAF1/C5 association did not correspond to any coding variant in either gene ^11^. TRAF1/C5 has also been linked by small studies to juvenile idiopathic arthritis (JIA), although the most extensive genetic study of JIA failed to detect a signal achieving genome-wide statistical significance ^12–15^.

*TRAF1* encodes tumor necrosis factor (TNF) receptor associated factor 1, expressed in activated myeloid and lymphoid cells ^16^. TRAF1 mediates pro-survival signaling downstream of TNF receptor superfamily members and negatively regulates Toll-like receptor and Nod-like receptor signaling ^17,18^. *C5* encodes complement factor 5, also implicated in arthritis biology by human and animal studies, rendering both genes plausible candidates ^11^. More recent functional data implicate *TRAF1* as the likely regulatory target since individuals with the high-risk haplotype exhibit lower TRAF1 expression and greater LPS-induced cytokine production in monocytes ^19^. However, the causal variant in TRAF1/C5 remains unknown, as is the pathway by which it reduces TRAF1 expression and whether the same mechanism is relevant for both RA and JIA.

Extending a prior ImmunoChip study of JIA in an expanded cohort of patients and controls, we confirm a disease association and prioritize a region within TRAF1/C5 as likely to contain the causal variant according to colocalization analysis. Using SNP-seq, bioinformatic prediction, and experimental validation, including donors bearing risk and protective alleles, we establish rs7034653 as a causal variant. We identify the transcription factor FRA2 as the regulatory protein, targeting this protein as orthogonal confirmation of the pathway. Conditioning on rs7034653 eliminates all genetic risk for RA at this locus, implicating a single shared pathway for both JIA and RA. Together, these findings define the mechanism underlying one of the first non-coding variants identified in inflammatory arthritis and model a strategy to transform GWAS findings into biologic understanding.

## Results

### Identification of rs7039505 as the lead SNP in the TRAF1/C5 locus in JIA

To evaluate whether genetic risk for JIA is conferred by the TRAF1/C5 locus, we studied genotype data from the ImmunoChip Consortium study of oligoarticular and seronegative polyarticular JIA, including from 2,756 published patients and 12,944 published controls together with an additional 1,183 patients and 1,468 controls ^20^. SNP rs7039505 achieved ImmunoChip significance (P=2.39 × 10^-7^, odds ratio=1.15, 95% confidence interval 1.09-1.21), confirming the presence of a causal variant in this locus (**Figure 1**). This SNP resided in close physical proximity but incomplete linkage disequilibrium (LD) with other variants associated previously with risk of JIA and RA (R^2^=0.74 for both, **Supplementary Figure 1A and 1B**). To assess whether the causal risk variant in the locus is the same variant that drives TRAF1 expression in monocytes, we performed colocalization analysis between the JIA ImmunoChip locus and eQTL datasets in monocytes and macrophages. We found high probability (PP4 > 0.8) that the causal variant for JIA colocalizes with the eQTL for TRAF1 expression in monocytes stimulated with LPS ^21,22^ as well as for other activated and resting states in monocyte and macrophages ^23,24^ (**Supplementary Figure 1C**). These studies indicate that the TRAF1/C5 locus likely contains a *TRAF1*-driving causal variant for JIA.

**Figure 1.**
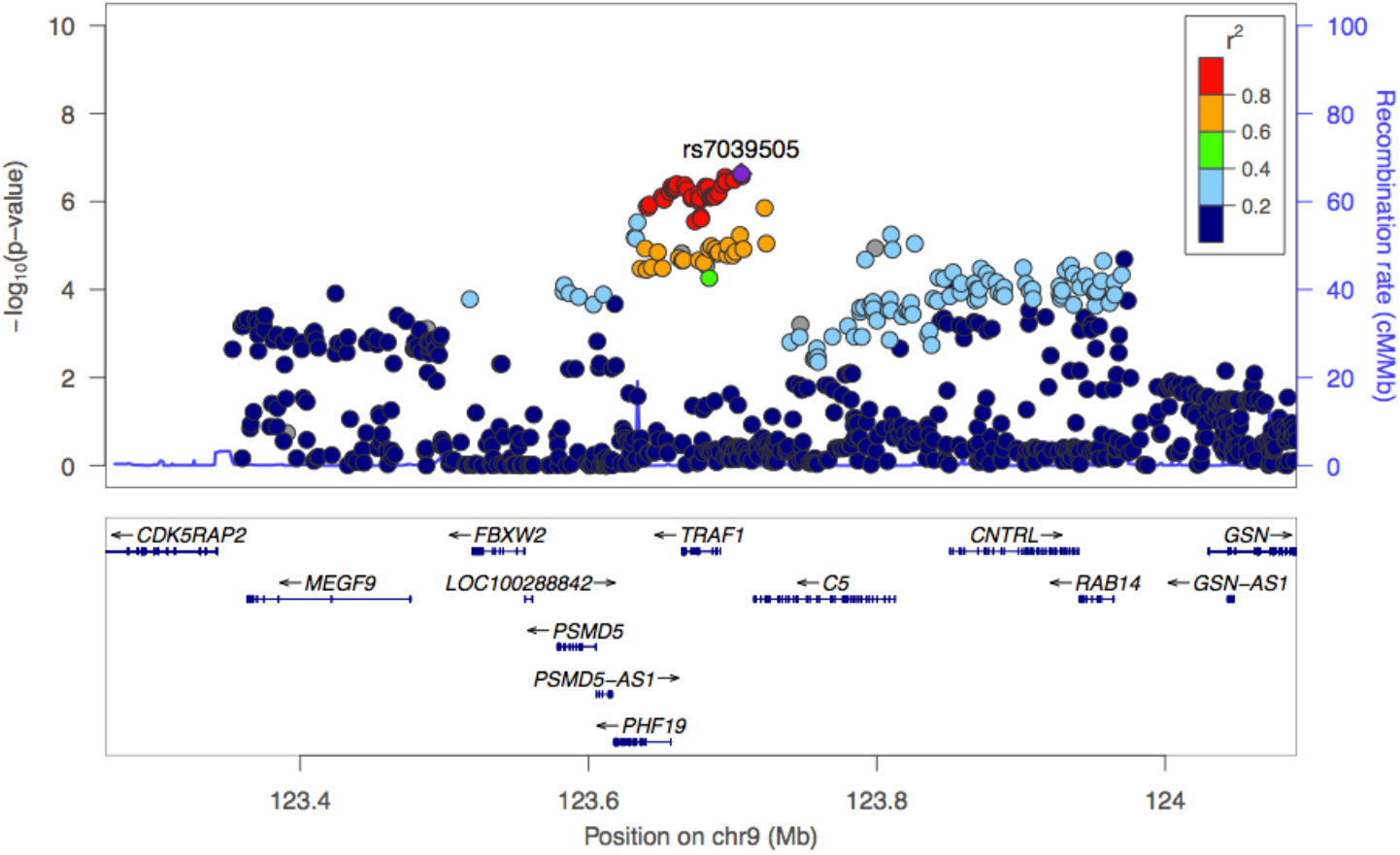
ImmunoChip analysis of the TRAF1/C5 locus in oligoarticular and seronegative polyarticular JIA. To define the non-coding variant within the TRAF1/C5 locus, we employed ImmunoChip data from 3,939 patients with oligoarticular and seronegative polyarticular JIA, and 14,412 controls. SNP rs7039505 achieved ImmunoChip significance (P=2.33 × 10^-7^,odds ratio=1.15, 95% confidence interval 1.09-1.21), confirming a genetic association of this locus.

### Identification of candidate causal variants via SNP-seq

To establish the basis of the association between rs7039505 and JIA, we identified all common variants in LD with this SNP from 1000 genomes Phase 3 (EUR population), filtered on minor allele frequency (MAF) >1%, LD R^2^>0.7, and location within a 1Mb region centered upon the tagging SNP. This strategy yielded 132 candidate SNPs for downstream screening and validation (**Supplementary Table 1**).

We then applied SNP-seq to narrow the candidate pool. In this method, a library of double-stranded DNA constructs is generated containing each allele of each candidate variant. A 31bp segment centered on the SNP of interest is flanked by binding sites for type IIS restriction enzymes (IISRE) that cleave DNA in a sequence-independent manner at 16bp (5’) adjacent to their binding site, i.e. at the SNP itself. Incubation of the library with nuclear extract followed by IISRE and next-generation sequencing (NGS) identifies variants protected from enzymatic restriction by transcription factors (TF) or other nuclear proteins (**Figure 2A**) ^2^. For this study, our library contained 271 dsDNA constructs reflecting each allele of each SNP, including 127 SNPs with two common alleles, 4 with three alleles, and 1 with five alleles (**Supplementary Table 3**). The library was incubated with nuclear extract from human monocyte-derived macrophages, washed, and exposed to the IISRE BpmI. As a control, the construct pool was incubated without nuclear extract. Surviving sequences were amplified by PCR and used as the input library for the next round of selection, repeating the procedure for 10 cycles, with barcoding at cycles 4, 6, 8, and 10 to identify over-represented SNPs. The entire procedure was performed in three independent replicates, with high correlation between sample pairs (Spearman ρ>0.9) (**Figure 2B**).

**Figure 2.**
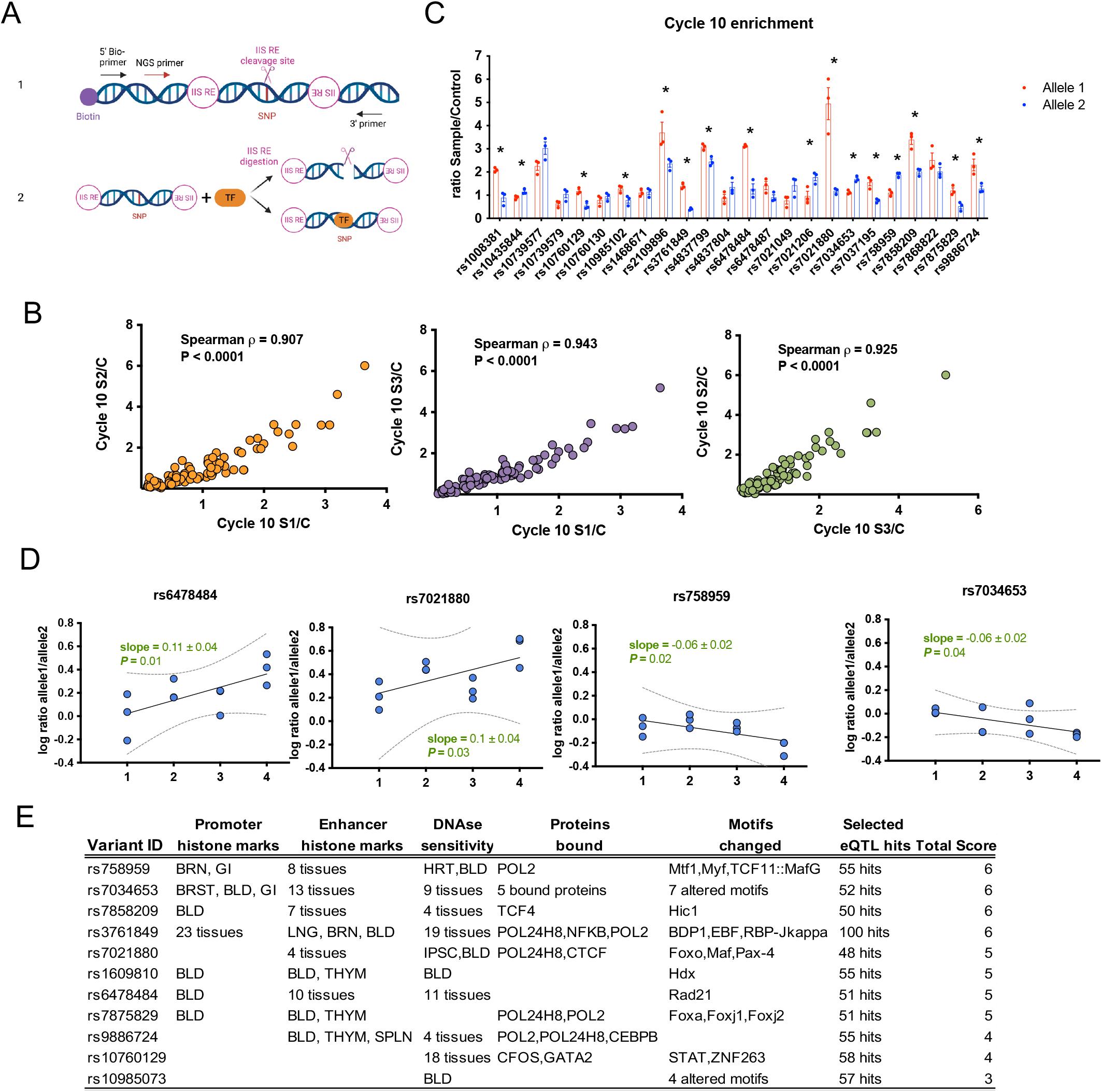
Screening candidate regulatory variants within TRAF1-C5 by SNP-seq. **(A)** SNP-seq (1) SNP-seq construct. A 31 bp sequence with the SNP centered in the middle on a type IIS restriction enzyme (IIS RE) cutting site is flanked with two IIS RE binding sites. A primer is included for high-throughput sequencing. (2) SNPs that fail to bind regulatory proteins such as transcription factors (TF) are negatively selected by PCR after type IIS RE cleavage (top right); protected SNPs can then be enriched by PCR (bottom right). The whole construct can be amplified using primers as per Supplementary Table 2. Bio, biotin; NGS, next generation sequencing. Created with BioRender.com **(B)** Correlation analysis of the NGS data for all three samples in SNP-seq, using nonlinear regression. (**C)** Reads count ratio between samples and controls at cycle 10 for 24 SNPs protected from Bpml cutting, Allele 1 and 2 for each SNP is shown in red and blue. 16 SNPs show significant allele specific protection. Statistical analysis was performed using multiple unpaired t test. (* represents p<0.05, n=3). (**D)** Top four SNPs showing an increment of allele-specific protection across cycles (cycle 4, 6, 8 and 10). A linear model for the ratio of reads in sample vs. control for allele 1 and allele 2, with an absolute slope greater than 0.05. P value is shown in each figure. (**E)** Annotations for the 11 candidate regulatory SNPs selected from SNP-seq using HaploReg.

We identified SNPs demonstrating allele-specific protection using two different NGS data standards, quality control (QC) 6 or QC12, matching sequencing data for either 6 or 12 nucleotides on either side of the SNP with the original sequences, and then employed two parallel analytical approaches established previously ^2^. First, we selected SNPs that exhibited a difference in protection between alleles of greater than 20% in cycle 10; of 24 SNPs proportionately enriched after SNP-seq compared with the input library, 16 exhibited such an allelic differential for each of the 2 QC standards (**Figure 2C**). Second, we sought SNPs with progressive allele-specific protection across cycles, fitting the ratio of reads for allele 1 and allele 2 in sample vs. control across cycles 4, 6, 8, and 10 and selecting SNPs with an absolute slope greater than 0.05; 10 SNPs exhibited this pattern for QC12 and 9 for QC6, including 4 that met both criteria for both standards (**Figure 2D, Supplementary Figure 2**). In the end, 11 SNPs that met both criteria from either QC standard were considered candidate causal SNPs, a list that did not include either the original tagging SNP rs7039505 or the lead tagging SNP for RA, rs3761847. All 11 candidates proved plausible based on proximity to epigenetic histone marks, DNAse hypersensitivity, sequence preference for TF binding, and association with gene expression as eQTL loci (**Figure 2E**).

### Identification of rs7034653 as the likely causal SNPs at *TRAF1-C5*

To confirm allele-specific protein binding, we employed the electrophoretic mobility shift assay (EMSA) using biotinylated DNA fragments of 31bp centered upon each allele of each candidate SNP, together with nuclear extracts from the monocytic cell line THP-1 and from human monocyte-derived macrophages. The SNPs rs7021880, rs7034653, rs9886724 and rs10985073 exhibited allele-imbalanced binding for both nuclear extracts, while rs758959 and rs1609810 exhibited this property for only THP-1 or only macrophage nuclear extract, respectively (**Figure 3A**, SNPs with allele-imbalanced nuclear protein binding in red, shifted bands marked as red dots). We then tested the allele-specific gene regulatory capacity of these 6 SNPs using a luciferase reporter. We cloned the 31bp DNA fragment containing either the risk or non-risk allele into the pGL3 vector and then transfected each reporter construct into THP-1, together with control vector pRL to normalize for transfection efficiency. We found that only rs7034653 and rs1609810 exhibited a significant difference between alleles (**Figure 3B**). Given the focus of prior work on rs3761847 ^19^, we evaluated the alleles of this SNP using both EMSA and the luciferase reporter assay. While the SNP exhibited allele-imbalanced binding in EMSA using both nuclear extracts, we observed no imbalanced luciferase reporter activity (**Figure 3C**).

**Figure 3.**
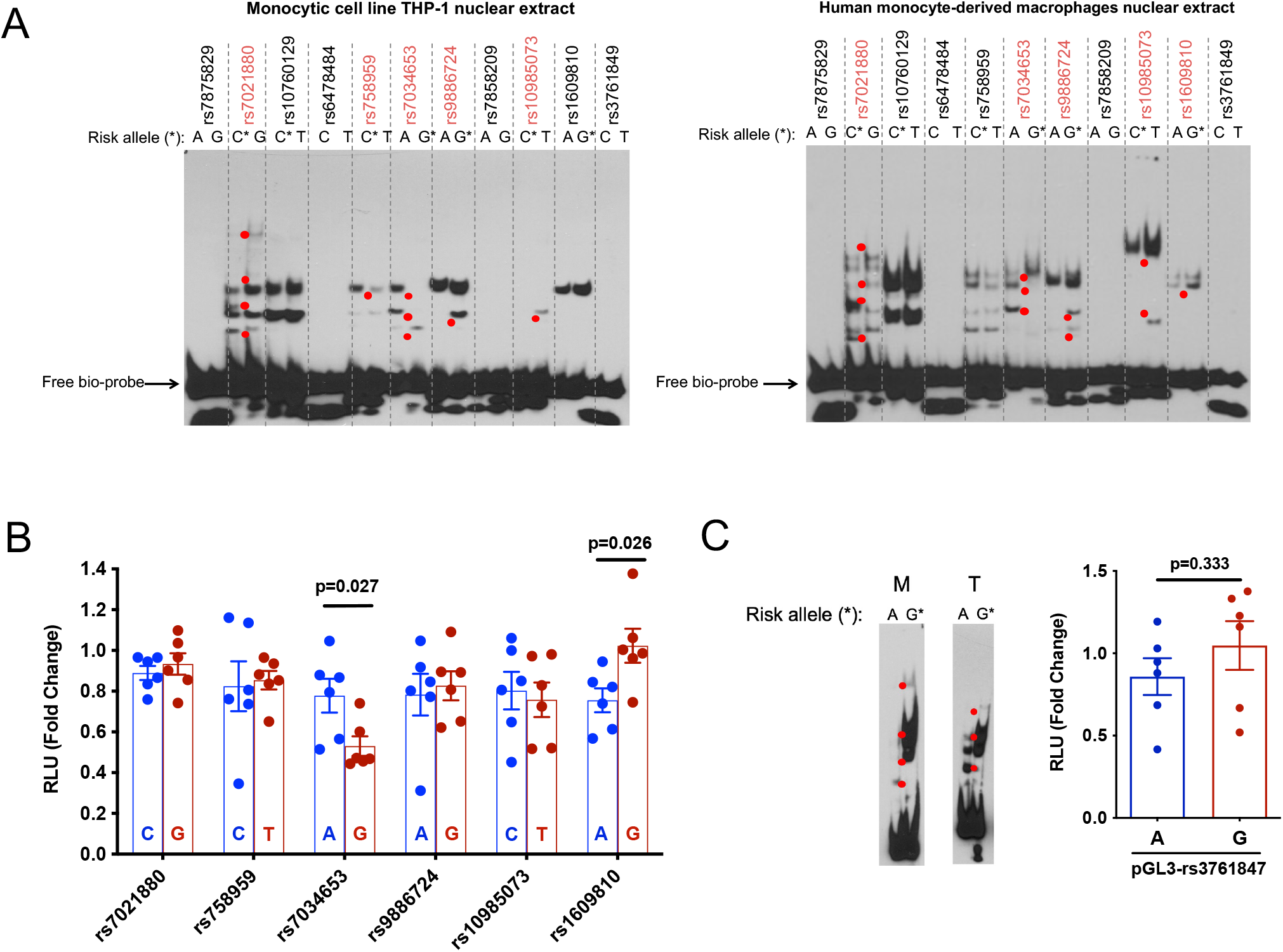
EMSA and luciferase reporter assessment of candidate regulatory SNPs. **(A)** EMSA for 11 candidate regulatory SNPs. Allele-specific gel shift/binding are shown as red dots. Left panel shows result with nuclear extract from monocytic cell line THP-1 cells and right panel shows result with nuclear extract from human monocyte-derived macrophages (*n* = 3 independent biological replicates with similar results). **(B)** Luciferase reporter assay showing relative luciferase activity in human THP-1 cells between the non-risk (blue) and risk (red) alleles of 6 candidate regulatory SNPs from EMSA. (mean ± s.d., *n* = 6 biological replicates, *t* test with two tails without correction for multiple hypothesis testing). **(C)** EMSA and luciferase reporter assay for SNP rs3761847. M indicates human monocyte-derived macrophages and T indicates THP-1 monocytic cell line. For luciferase assays, histogram shows mean ± s.d., *n* = 6 biological replicates, *t* test with two tails without correction for multiple hypothesis testing. P values for figure B and C are shown in the figures.

We then employed publicly available data to further assess rs7034653 and rs1609810. Using DICE (Database of Immune Cell Expression, eQTLs, and Epigenomics), we found that both SNPs were associated with expression of *TRAF1* as well as of the noncoding transcript *PSMD5-AS1*. Since *PSMD5-AS1* has no known function, we focused on *TRAF1*, seeking to identify an eQTL pattern consistent with our experimental data. Per DICE, the G allele of each SNP is associated with lower *TRAF1* expression in monocytes (**Supplementary Figure 3A**), consistent with our luciferase result for rs7034653 but opposite that observed for rs1609810. The BIOS QTL browser database also revealed rs7034653 as the best candidate to regulate *TRAF1* expression based on methylation quantitative trait loci (**Supplementary Figure 3B**). Finally, using a bioinformatic model developed to identify sequence-specific binding proteins ^25,26^, we found that the likely high-binding allele for these SNPs was the A allele for both rs7034653 (top binding candidate TF, MAF BZIP Transcription Factor G, MAFG) and rs1609810 (top binding candidate TF, Highly Divergent Homeobox) (**Supplementary Table 4**); since only the A allele of rs7034653 exhibited preferential gel shift in EMSA, again rs7034653 appeared the more likely candidate. Considering these data together, we selected rs7034653 and its risk allele G (MAF 32% from TopMed) for further functional study as the best candidate regulatory SNP at TRAF1/C5.

### rs7034653 risk allele G is associated with lower TRAF1 expression and increased pro-inflammatory response after LPS stimulus

The arthritis risk haplotype at TRAF1/C5 is believed to reduce expression of *TRAF1* and thereby enhance production of inflammatory mediators ^19^. We therefore tested whether individuals differing in allele carriage at rs7034653 exhibited the predicted variation in TRAF1 expression and TNF release. We recruited healthy donors genotyped through the Mass General Brigham Biobank using Joint Biology Consortium (JBC) infrastructure (www.jbcwebportal.org). We isolated peripheral blood monocytes by negative selection from 13 donors of AA, AG and 12 donors of GG genotypes at rs7034653. TRAF1 is expressed at low levels in resting monocytes but induced via NFκB, so we selected LPS as the appropriate stimulus. At baseline, TRAF1 expression was similarly low in all groups; however, after LPS exposure, expression of TRAF1 increased more in the AA (protective) than GG (risk) monocytes, with AG cells exhibiting an intermediate phenotype (**Figure 4A, 4B**; gating strategy for CD14+HLA-DR+ monocytes shown in **Supplementary Figure 4A**). Correspondingly, LPS-stimulated TNF release was greatest for GG monocytes (**Figure 4C**). Intracellular staining for TNF, IL-6 and IL-1β as well as IL-6 release were more variable, such that we could not observe significant differences by genotype, although GG monocytes trended higher across most measures (**Supplementary Figure 4B, 4C**).

**Figure 4.**
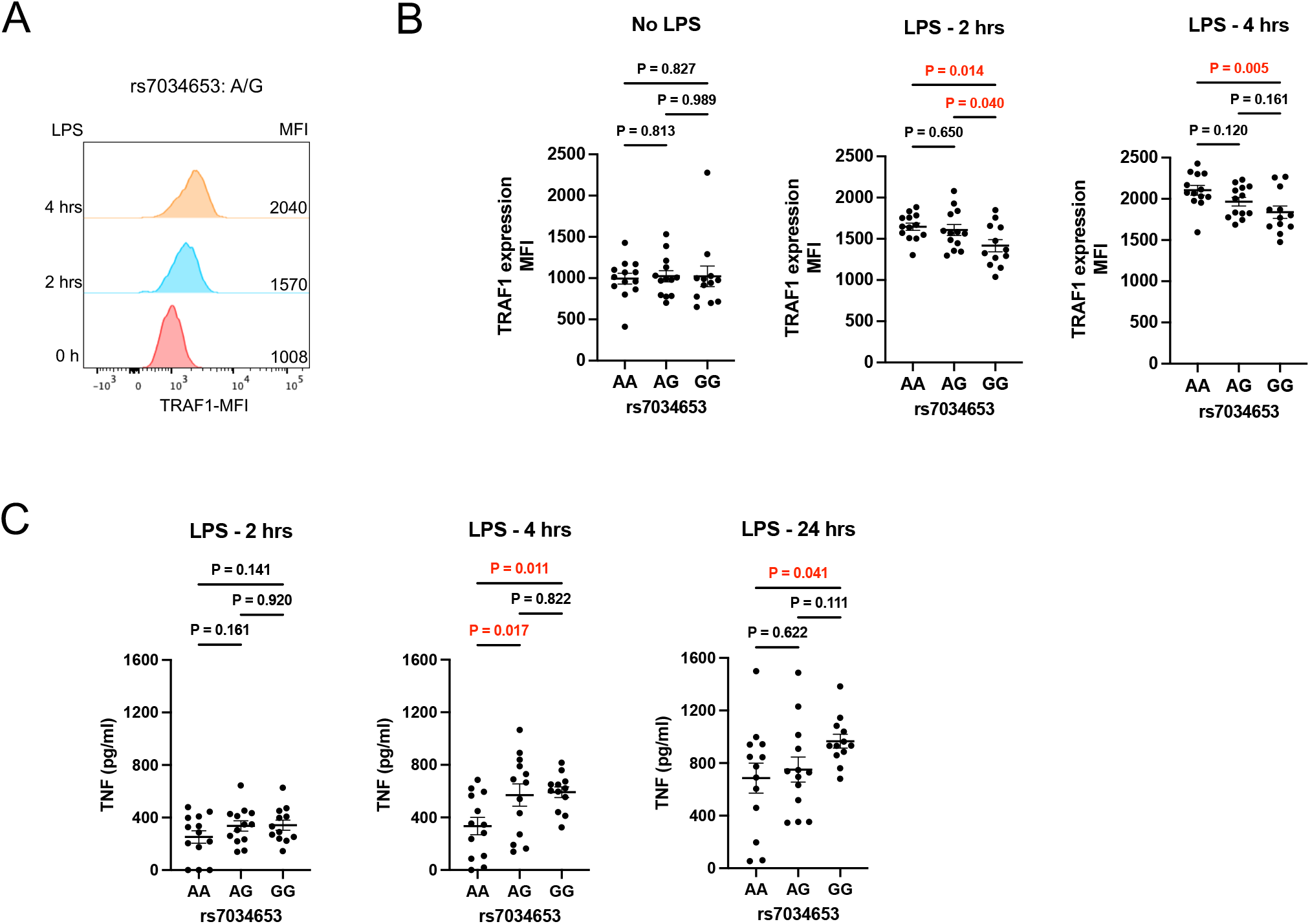
Primary human monocytes bearing the GG risk haplotype at rs7034653 express lower TRAF1 and produce more inflammatory cytokines. Purified monocytes from healthy human donors were treated or not with LPS (100 ng/ml for 2 or 4 hours), and TRAF1 protein was measured by flow cytometry, TNF protein was measured by ELISA. **(A)** Histograms showing the time course of induction of TRAF1 by LPS from purified monocytes from one representative donor (AG genotype at rs7034653). **(B)**. Mean fluorescence intensities (MFI) of TRAF1 in purified monocytes of subjects with the AA, AG and GG genotypes at rs7034653 at different time points. **(C)** ELISA measurements of TNF in monocyte supernatants after LPS stimulation. Each symbol represents one donor; n = 13, 13 and 12 human subjects of genotypes AA, AG and GG, respectively. Statistical analysis was performed using one-way ANOVA with multiple comparisons. All p values are shown in the figure.

### FRA2 is the transcription factor that binds rs7034653 to regulate TRAF1

rs7034653 is located in the third intron of *TRAF1*, suggesting that its impact on TRAF1 expression is likely mediated through binding of a TF, a conclusion supported by our SNP-seq, EMSA, and luciferase results. To provide an orthogonal line of evidence implicating rs7034653 in the arthritis risk conferred through the TRAF1/C5 locus, we sought to identify the TF and thus the regulatory pathway mediating enhanced pro-inflammatory mediator production. We employed a bioinformatic approach to identify TFs likely to bind this SNP based on protein-binding motif preferences ^25,26^. Several candidate TFs were identified as having the potential to bind rs7034653 in an allele-imbalanced manner (**Supplementary Table 4**). Given the findings from EMSA that the causal TF likely exhibited preferential binding to the A allele, we performed oligo pull down with this allele using nuclear extract from THP-1 cells, with or without excess non-biotinylated competitor and employing the incidental TRAF1/C5 SNP rs1609810 as a control. Among the candidate TFs, Fos-Related Antigen 2 (FRA2), Growth Factor Independent 1 Transcriptional Repressor (GFI1), and Activating Transcription Factor 3 (ATF3), only FRA2 exhibited the expected specific binding to rs7034653 (**Figure 5A**). This result was confirmed by EMSA-supershift, finding that binding to rs7034653 was weakened by anti-FRA2 antibody (disappearance of the band) compared with isotype control or antibody against another TF candidate, MAFG (**Figure 5B**). These studies establish that FRA2 can bind the rs7034653 sequence.

**Figure 5.**
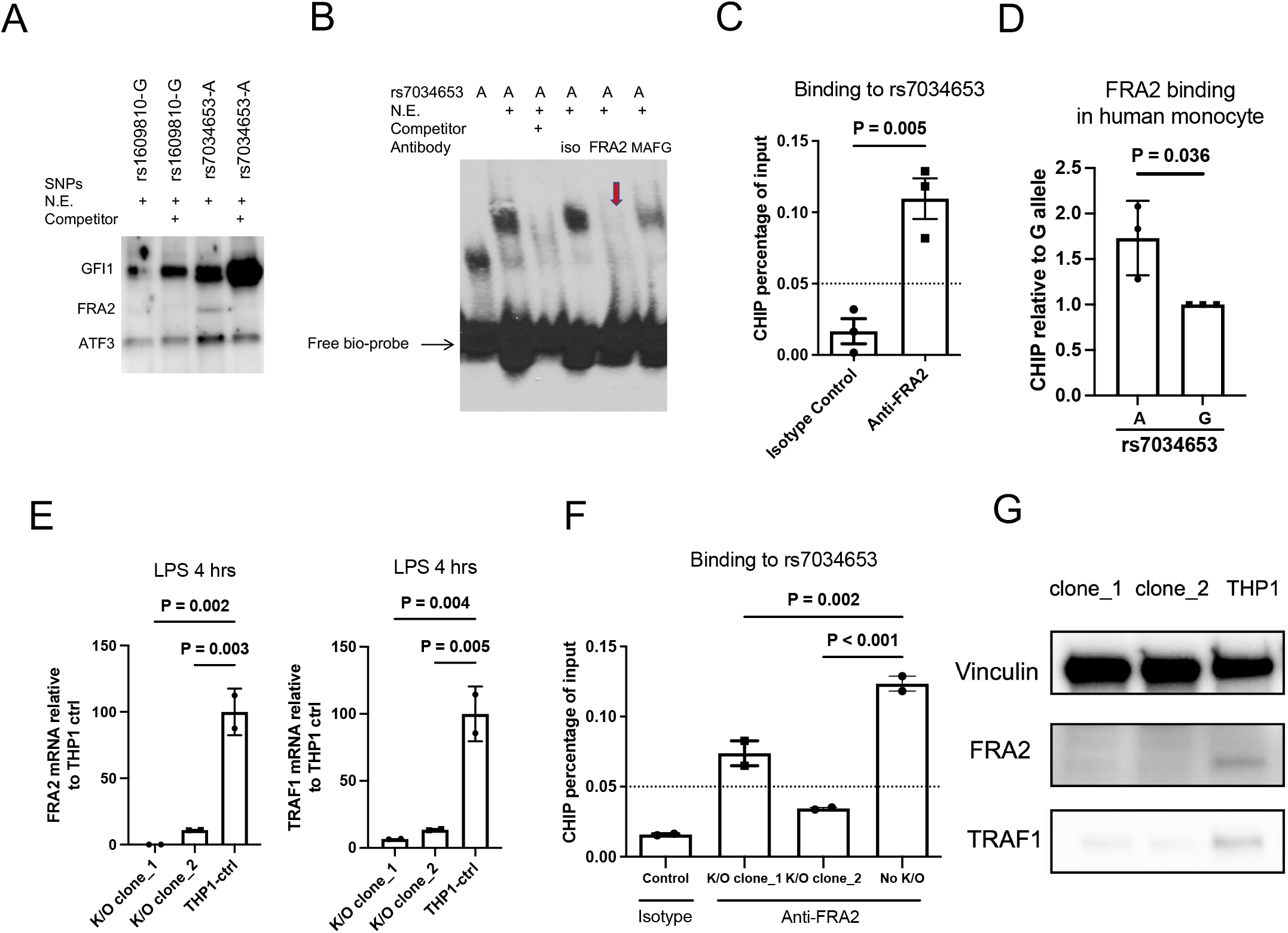
FRA2 binding to rs7034653 regulates TRAF1 expression. **(A)** Oligonucleotide pulldown western blot assay shows FRA2 binds specifically to rs7034653-A; binding is eliminated by 30-fold excess of non-biotinylated rs7034653-A competitor. GFI1 and ATF3 bind to all samples with or without non-biotinylated competitors. **(B)** EMSA supershift assay shows specific binding of rs7034653-A to FRA2. The addition of antibody to FRA2 in the oligo/nuclear extract (THP-1) mixture caused the disappearance of shift band (arrow), whereas isotype control antibody and antibody to irrelevant control MAFG did not. **(C)** CHIP-qPCR shows FRA2 binding to rs7034653 in human THP-1 cells (mean ± s.d., *n* = 3). **(D)** Allele discrimination CHIP-qPCR shows FRA2 preferentially binds to A allele over G allele of rs7034653 in unstimulated human monocytes heterozygous at rs7034653. (mean ± s.d., *n* = 3). **(E, G)** CRISPR-Cas9 FRA2 knockout THP-1 clones show decreased FRA2 and TRAF1 mRNA and protein expression compared with negative sgRNA-treated THP-1 control. (mean ± s.d., *n* = 2). **(F)** CRISPR-Cas9 FRA2 knockout THP-1 clones show decreased FRA2 binding to rs7034653 compare with negative sgRNA-treated THP-1 control. (mean ± s.d., *n* = 2). Panels A, B, E, F, G representative of duplicate biological replicates, panels C and D, triplicate biological replicates. Statistical analysis methods used for panels C and D were un-paired *t* test with two tails without correction, panel E and F were one-way ANOVA with multiple comparisons. All p values are shown in the figure.

To confirm *in vivo* interaction between rs7034653 and FRA2, we performed chromatin immunoprecipitation (ChIP)-qPCR. Anti-FRA2 antibody was able to pull down rs7034653 significantly more than isotype control antibody in THP-1 cells (AA for rs7034653) (**Figure 5C**). In purified human monocytes from heterozygous donors, we observed more efficient pulldown of the A allele than the G allele by anti-FRA2, consistent with our bioinformatic and experimental predictions (**Figure 5D**).

To test whether the interaction of FRA2 with rs7034653 modulated *TRAF1* expression, we generated two *FRA2* knockout clones using CRISPR-Cas9. Both clones showed *FRA2* deficiency and reduced *TRAF1* expression after LPS stimulation (**Figure 5E**). As expected, deficiency of FRA2 markedly attenuated the ability of anti-FRA2 to pull down rs7034653 via ChIP (**Figure 5F**). Gene deletion translated into significantly decreased protein expression of FRA2 and, in parallel, TRAF1 (**Figure 5G**), confirming the role of this SNP/TF interaction in TRAF1 expression.

### Relationship between rs7034653 and RA risk in the TRAF1/C5 locus

Finally, we sought to determine whether the causal SNP identified here could account for some or all of the GWAS signal at this locus for RA ^11^. Until recently, the lead tagging SNP for RA was rs3761847, which resides in close physical proximity but imperfect LD with rs7034653 (R^2^=0.77, p<0.0001) ^11^. However, an expanded trans-ancestry GWAS with improved fine-mapping resolution and higher statistical power has now identified rs1953126 as the lead SNP at TRAF1/C5 (p=3.18 × 10^-11^, **Figure 6A**) ^10^. This SNP exhibits much higher LD with rs7034653 (R^2^=0.98, p<0.0001). Conditioning on rs7034653 in 25 European RA GWAS datasets resolved all risk signal (**Figure 6B, C**). This finding shows that rs7034653 accounts for all locus-associated disease risk in RA, implicating a shared mechanism at TRAF1/C5 in in RA and JIA.

**Figure 6.**
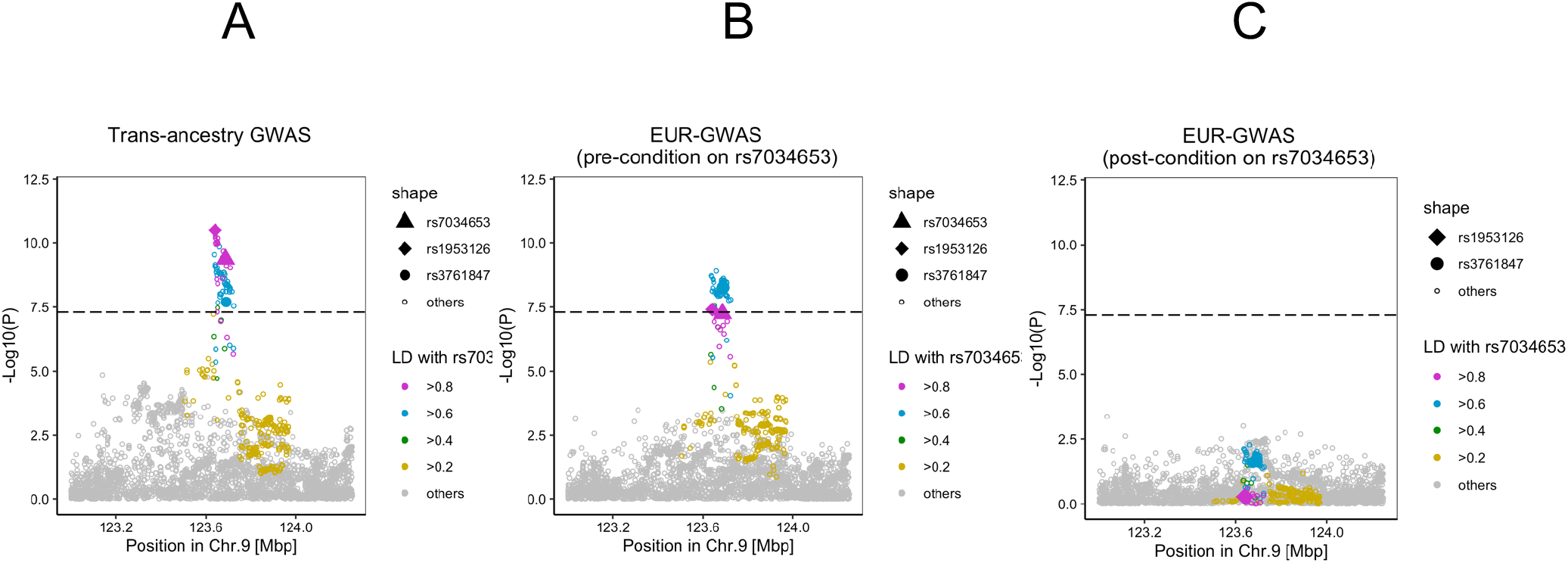
Conditional analysis on rs7034653 confirms no residual risk in rheumatoid arthritis. (A) TRAF1/C5 locuszoom plot in trans-ancestry GWAS of RA. (B) Pre-conditional analysis locuszoom plot in TRAF1/C5 in the European-ancestry cohort, comparable to the JIA ImmunoChip population. (C) Post-conditional analysis in the European population detected no residual RA risk. Conditional analysis was conducted in each cohort of the 25 European-ancestry GWAS and the results were meta-analyzed using the inverse-variance weighted fixed effect model.

## Discussion

Inflammatory arthritis encompasses a range of phenotypes, historically dichotomized by age of onset into JIA and RA ^27^. Here, beginning from an ImmunoChip study of oligoarticular and seronegative polyarticular JIA, we identified the non-coding variant rs7034653 as a causal variant within TRAF1/C5 that mediates risk of both JIA and RA through differential allelic affinity for the transcription factor FRA2. Individuals bearing the G risk allele bind FRA2 less strongly than those with the A allele, resulting in attenuated upregulation of the anti-inflammatory protein TRAF1 and thereby enhanced production of TNF from stimulated monocytes (**Figure 7**). These findings inform the mechanism through which one of the earliest GWAS “hits” functions in arthritis and model a general approach to the investigation of noncoding causal variants in human polygenic disease.

**Figure 7.**
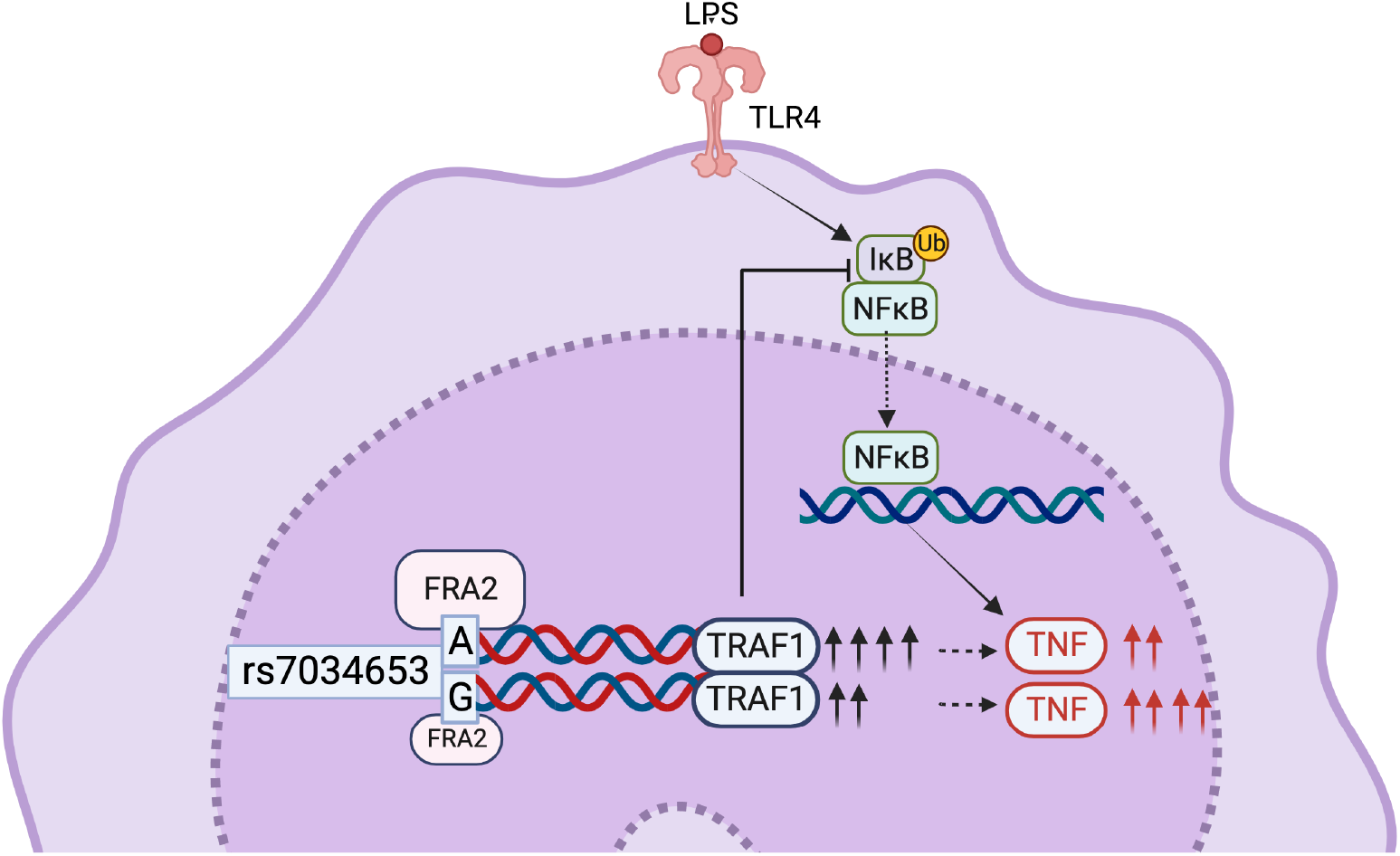
Allelic variation at rs7034653 modulates TNF production through differential binding of FRA2 to regulate expression of TRAF1. FRA2 exhibits greater affinity for the protective A allele than the risk G allele of rs7034653, leading to enhanced production of the negative regulatory protein TRAF1 and lower production of the pro-inflammatory mediator TNF. Created with BioRender.com.

FRA2 is a member of the activator protein 1 (AP-1) family, whose roles involve regulating gene expression in response to a variety of stimuli, including cytokines, growth factors, stress, and bacterial and viral infections ^28^. Encoded by *FOSL2*, FRA2 forms a heterodimer with other AP-1 members to regulate downstream gene expression ^28^. Expression of FRA2 is induced by LPS but also by cytokines such as IL-1 and TNF, mediators abundant in the inflamed joint and implicated in arthritis pathogenesis both by animal models and by the therapeutic efficacy of cytokine blockade in human patients ^29–35^. Concordant with earlier investigations of the TRAF1/C5 locus ^19^, our findings suggest that impaired induction of the counterregulatory factor TRAF1 shifts the balance toward amplified rather than suppressed inflammation to confer excess risk of inflammatory arthritis.

It is notable that many variants at TRAF1/C5 exhibited allele-specific differences in SNP-seq, EMSA, and luciferase steps, as well as via bioinformatic plausibility filters, and it was only by combining these that we could narrow the candidate pool to a single SNP. These results illustrate the complexity intrinsic to experimental study of non-coding variants, and the need for additional complementary approaches – in this case identification of the causal transcription factor – to provide confidence in one over others. Our data do not exclude the possibility that the TRAF1/C5 risk haplotype harbors more than one causal variant. For example, studies of *STAT4* identified 2 protein-binding SNPs, tightly linked within one haplotype, that each bound distinct transcription factors regulating the gene ^2^. However, the most parsimonious explanation for our findings is that rs7034653 drives TRAF1/C5-associated disease risk within the population studied.

Genetic variants can exhibit differential effects on cell function depending on lineage and activation state ^36^. Both TRAF1 and FRA2 are expressed widely across immune cell types. We selected the monocyte/macrophage lineage for our SNP-seq screen and downstream validation studies, but it is possible that effects in other cell types, or mechanisms other than de-repressed cytokine production, contribute to arthritis risk. For example, TRAF1 forms a dimer with TRAF2 to transduce signaling from TNF and related molecules to suppress TNF-induced apoptosis ^16,18,37^. Correspondingly, mice lacking TRAF1 are highly susceptible to both LPS-induced septic shock and TNF-induced skin necrosis ^18,19^. In T cells, TRAF1 deficiency enhances susceptibility to TNF-induced activation and proliferation ^17,18^. In B cells, TRAF1 cooperates with TRAF2 in induction of NF-kB via CD40 ^38^, while B cell chronic lymphocytic leukemia cells show overexpression of TRAF1 ^17^. TRAF1 also enhances classical signaling by NF-κB and mitogen associated protein (MAP) kinases downstream of TNFR to promote lymphocyte survival ^37–41^. Thus, differential genetic control of TRAF1 by FRA2 could have multiple ramifications for immune-mediated disease, although enhanced macrophage-derived cytokine production remains a plausible mechanism given the transformative therapeutic impact of TNF blockade in both JIA and RA.

By implicating a shared variant in TRAF1/C5 in both JIA and RA, our findings support the close relationship between these disease families. The sharp division in arthritis nomenclature between childhood and adult arthritis arose in the 1950s out of descriptive convenience, reflecting the age of patients cared for in pediatric centers rather than any established biological differences ^27^. Our ImmunoChip data are derived from children with either oligoarticular or polyarticular onset who lack circulating rheumatoid factor, a population that includes three subgroups in the current JIA nomenclature: persistent oligoarticular JIA, extended oligoarticular JIA, and seronegative polyarticular JIA. These patients share HLA and non-HLA associations with adult seronegative arthritis ^27,42–44^. Interestingly, TRAF1/C5 was originally identified in an RA population composed primarily of seropositive patients, a group that also transcends the pediatric/adult boundary but that exhibits different HLA associations compared with seronegative disease ^11,45–47^. The fact that the rs7034653/FRA2/TRAF1 mechanism confers risk for both seronegative and seropositive arthritis suggests a role in a shared downstream mechanism, such as synovial inflammation. This possibility is further supported by the fact that the TRAF1/C5 locus is not a GWAS “hit” for autoimmune diseases such as type I diabetes and autoimmune thyroid disease, underscoring the potential relevance of excess mediator production by myeloid cells as a key mechanism by which rs7034653 predisposes to arthritis.

Our findings have clinical implications. Identification of a specific SNP as causal enables greater ability to test genetic variation as a tool for patient stratification. We identify the FRA2/TRAF1 pathway as a relevant target across the arthritis spectrum, raising the possibility that enhancing expression of TRAF1, or controlling TRAF1-regulated pathways, could yield important therapeutic benefit. More broadly, the approach shown here for TRAF1/C5 could be employed across loci and across diseases to define the mechanistic implications of the regulatory variants that are thought to underlie most GWAS “hits”, helping to translate the major public investment in genetic studies of complex polygenic disease into tangible advances in pathogenic understanding and treatment.

## Author Contributions

QW, MM-B, PAN conceived and designed the study. QW, M-MB, TK performed experiments and performed data analysis. JAS led recruitment of genotyped human subjects. KI, MS, SP, SR, CDL, MCM, SDT provided and analyzed genetic data. VA, MCH, MG-A, XTC, MTW provided data analysis. QW, PAN drafted the manuscript and all authors edited and approved the manuscript. PAN supervised the research.

## Acknowledgements

QW, M-MB, and TK were supported by Joint Biology Consortium microgrants off parent grant NIH/NIAMS P30AR070253. JAS was supported by NIH/NIAMS R01 AR077607, P30 AR070253, and P30 AR072577), the R. Bruce and Joan M. Mickey Research Scholar Fund, and the Llura Gund Award for Rheumatoid Arthritis Research and Care. MTW was supported by NIH grants R01NS099068, R01HG010730, U01AI130830, R01AI024717, R01AR073228. MS was supported by P30AR070549. VA, MCH, M-GA were supported by NIH/NIAMS P30AR069625, Gilead Sciences, Lupus Research Alliance, and the Arthritis National Research Foundation Vic Braden Family Fellowship. SP was supported in part by the Marcus Foundation Inc., Atlanta, GA. SR was supported by 5R01AR063759-07. CDL and MCM were supported by R01 HD089928-05. SDT was supported by P30AR070549. PAN was supported by NIH/NIAMS 2R01AR065538, R01AR075906, R01AR073201, P30AR070253, the Fundación Bechara, and the Arbuckle Family Fund for Arthritis Research. The Trøndelag Health Study (HUNT), which contributed with control samples, is a collaboration between the HUNT Research Centre (Faculty of Medicine and Health Sciences, NTNU - Norwegian University of Science and Technology), Trøndelag County Council, Central Norway Regional Health Authority, and the Norwegian Institute of Public Health. We acknowledge Nils Thomas Songstad and Nina Moe for patient recruitment in the Norwegian subcohort of the Nordic JIA study, Kristin Rian, NTNU, Trondheim, for technical support, and Helse Nord Research Grants for funding collection of samples used in this study from Tromsø, Norway.

## Online Methods

### Genetic association study

Samples are from the ImmunoChip Consortium study of oligoarticular and seronegative polyarticular JIA, including 2,756 patients and 12,944 controls already published ^20^ together with an additional 1,183 patients and 1,468 controls (unpublished), totaling 3,939 JIA patients and 14,412 controls. Samples were genotyped using the Immunochip, a custom Illumina Infinium array, according to Illumina’s protocols at laboratories in Hinxton, UK; Manchester, UK; Cincinnati, USA; Utah, USA; Charlottesville, USA; and New York, USA. The Illumina GenomeStudio GenTrain2.0 algorithm was used to recluster all 18,351 samples. Single-nucleotide polymorphisms (SNPs) were initially excluded if they had a call rate <98% and a cluster separation score of <0.4. A SNP was subsequently removed from the primary analysis if it exhibited significant differential missingness between cases and controls (p<0.05), had significant departure from Hardy-Weinberg equilibrium (p<0.000001 in cases or p<0.01 in controls), or had a minor allele frequency (MAF) <0.01. Then, using only the SNPs that passed the above quality control thresholds, samples were excluded for any of the following: call rate <98%, inconsistencies between reported and genotype-inferred gender, or excess heterozygosity on the autosomes. Duplicates, first-degree, and second-degree relatives were identified using identity-by-descent statistics computed using the program KING ^48^. From these pairs, the sample with the highest call rate was retained. Admixture estimates were computed on the remaining samples while including the HapMap phase III individuals (CEU, YRI and CHB) as reference populations using the software ADMIXTURE ^49^. The admixture estimates were then used to identify and remove genetic outliers and including in the statistical models as covariates. A P value smaller than 1 × 10^-6^ was considered ImmunoChip significant ^50^.

### Colocalization

We performed colocalization analysis for the ImmunoChip variants in the TRAF1/C5 locus against the eQTL Catalogue ^51^. We imported gene expression QTL summary statistics from RNA-seq and microarray studies on naïve and stimulated monocytes or macrophages ^21–24,52–54^. We fetched the summary statistics data using the tabix method with the seqminer R package (v8.5). We included only biallelic SNPs and performed the analysis for genes with a transcription start site within a window of ±250,000 base pairs from the ImmunoChip lead variant, and for which there was at least one eQTL passing the 5 × 10^-5^ p-value threshold. Before merging ImmunoChip and QTL data, the variant coordinates of the ImmunoChip were lifted to the GRCh38 version of the reference genome using liftOver with the UCSC chain file. We used the coloc v5.1.0.1 package ^55^ in R v4.2.0 to test for colocalization at each gene and dataset. The code is available at <https://github.com/gutierrez-arcelus-lab/jia_coloc>.

### Cells and culture

Human monocytic cell line THP1 was purchased from ATCC (catalog number TIB-202) and cultured in RPMI1640 medium (ThermoFisher Scientific, catalog number 11875093) supplemented with 10% Fetal Bovine Serum (FBS) (ThermoFisher Scientific, catalog number 26140079) and 1% Penicillin-Streptomycin (ThermoFisher Scientific, catalog number 15140122), 0.05 mM 2-Mercaptoethanol (ThermoFisher Scientific, catalog number 21985-023), 1 mM Sodium Pyruvate (ThermoFisher Scientific, catalog number 11360070), 10 mM HEPES buffer (ThermoFisher Scientific catalog number 15630130).

### Human PBMC isolation and monocyte purification

For studies in genotyped donors, whole blood was collected from genotyped healthy human subjects recruited from the Mass General Brigham Biobank through Joint Biology Consortium (JBC) recruitment core (www.jbcwebportal.org). Genotype was confirmed in each donor by sanger sequencing. Peripheral blood mononuclear cells (PBMCs) were isolated from whole blood collected in BD Vacutainer™ Plastic Blood Collection Tubes with K2EDTA via Ficoll gradient (Cytiva, catalog number GE17-1440-02) and stored in liquid nitrogen in 10% DMSO in complete FBS. After thawing, CD14+ monocytes were isolated using the EasySep Human Monocyte Isolation Kit (Stemcell Technologies, catalog number 19359) according to manufacturer’s instructions. In other experiments, PBMCs were extracted from apheresis leukoreduction collar blood from the Brigham and Women’s blood donation center. Human monocyte derived macrophages were generated by culturing human monocyte purified from collar PBMCs in the presence of 15 ng/ml granulocyte-macrophage colony-stimulating factor (GM-CSF) (PeproTech, catalog number 30003) for 5 days in complete RPMI 1640 medium. Human subjects research with these samples was approved by the Brigham and Women’s Hospital IRB (protocol reference number 2019P003709).

### SNP-seq oligos and primers

All primers were purchased from IDT as listed in supplementary table 2. SNP-seq oligo pool was purchased from TwistBiosciences and oligos for EMSA screening and luciferase reporter assay was purchased from EtonBiosciences.

### SNP-seq

SNP-seq constructs were built according to Figure 2. The SNP sequence is 31 bp long, centered on the SNP of interest. The sequences of PCR amplification primers for the library oligonucleotides are listed in supplementary table 1. For oligo library amplification, 100 ng of pooled DNA was amplified with bioMagF-G5 and MagR-G3 by PCR for 25 cycles with AccuPrime Taq (Thermo Fisher Scientific, catalog number 12346086) at 94 °C for 60 s, 58 °C for 60 s and 68 °C for 40 s. After gel purification, 10 ng of biotinylated DNA was attached to 4 μl streptavidin-Dynabeads (Invitrogen, catalog number 11205D) according to the manufacturer’s protocol. The DNA-bound beads were then incubated with 60 μg nuclear extract from human monocyte derived macrophages or no nuclear extract for 1 h at room temperature in LightShift Chemiluminescent EMSA Kit reaction buffer (Thermo Fisher Scientific, catalog number 20148X). After washing and separation, the DNA-bound beads were digested with 2 μl BpmI (NEB, catalog number R0565L) for 30 min at 37 °C. After another wash and separation, the DNA was amplified again with bioMagF and MagR and reattached to the Dynabeads for the next SNP-seq cycle. Ten cycles were performed in total. DNA from cycle 4, 6, 8 and 10 were used to prepared Next Generation Sequencing (NGS) library by two consecutive PCRs (PCR with L1seq and R1 primers, 20 cycles of 98 °C 1:30 min, 60 °C 1:40 min, 72 °C 0:40 min; PCR with L2 and R2 primers, 25 cycles of 98 °C 1:30 min, 60 °C 1:40 min, 72 °C 0:40 min) for the independent samples using Herculase polymerase (Agilent, catalog number 600677). The primers used are also listed in supplementary table 1. PCR products were run in a 2% agarose gel and the correct band (around 200 bp) was purified with QiaQuick gel extraction Kit (QIAGEN, catalog number 28706) (at least 2 columns) following manufacturer’s instructions. The elution was sent for NGS at Dana Farber Genomics Core, Harvard Medical School.

### Electrophoretic mobility shift assay (EMSA)

EMSA was performed using the LightShift Chemiluminescent EMSA Kit (Thermo Scientific, catalog number 20148) according to manufacturer’s instructions. For oligo probe, a 31 bp fragment with the SNP centered in the middle was made by annealing two biotinylated oligonucleotides. Nuclear proteins were extracted from either THP-1 cells or human monocyte-derived macrophages using NE-PER Nuclear and Cytoplasmic Extraction Reagents (Thermo Scientific, catalog number 78835) per manufacturer instructions. For gel supershifts, 4 μg of antibody was added after for additional 30 min incubation. Antibodies used are anti-MAFG antibody (GeneTex, catalog number GTX114541), anti-FRA2 antibody (Abcam, catalog number ab124830) and IgG isotype control antibody (Thermo Scientific, catalog number 02-6102).

### Luciferase reporter assay

The luciferase reporter assay was performed exactly according to the manufacture’s manual (Promega, catalog number E1751). The pGL3 expression vector (0.35 μg) with or without SNPs sequence was co-transfected with the control vector pRLTK (0.25 μg) using TransIT-2020 (Mirus, catalog number MIR 5404) into 1 × 10^5^ THP-1 cells. After24 h incubation, luciferase activity was measured with the Dual-Glo luciferase assay system (Promega, catalog number E2920).

### Transcriptional factors binding prediction

Candidate TFs binding to rs7034653 was analyzed using a published model to identify sequence-specific binding proteins ^25,26^.

### Flow cytometry

For monocyte staining, cells were treated with LPS for the indicated times and then blocked with Fc Receptor Binding Inhibitor Polyclonal Antibody (eBioscience, catalog number 14-9161-71) and stained with live/dead-e506 stain (eBioscience, 65-0866-14), followed by staining with anti-HLA-DR–FITC (clone L243), anti-CD14–APC eFluor780 (clone 61D3) and anti-TRAF1-PE for intracellular TRAF1 staining or anti-HLA-DR–FITC (clone L243), anti-CD14–APC eFluor780 (clone 61D3), anti-IL-1β–PE (clone CRM-56), anti-IL-6–e450 (clone MQ2-13A5), anti-TNF–PerCP-Cy5.5 (clone MAb11) for intracellular cytokine staining; all the above antibodies were purchased from eBioscience or labeled manually, used at 1:200 dilution. Purified TRAF1 antibody (Millipore Sigma, catalog number MABC260) was labeled with PE (Biotium, catalog number 92299). For intracellular staining, cells were fixed with Foxp3/ Transcription Factor Staining Buffer Set (eBioscience, catalog number 00-5523-00) after surface marker staining. For intracellular cytokine staining, purified monocytes were treated with LPS for the indicated times and GolgiPlug (brefeldin A; BD Biosciences, catalog number 00-4506-51) was added in the final 4 h of LPS stimulation. For intracellular TRAF1 staining, supernatants were collected from each sample and saved for cytokine ELISA measurement. Cells were analyzed on a BD FACSCanto II (BD Biosciences).

### ELISA

human TNF (Thermo Scientific, catalog number 88-7346-22) and IL-6 (Thermo Scientific, catalog number 88-7066-22) ELISA were performed according to manufacturer’s instructions.

### Pulldown assay

Pulldown assay is modified from a published protocol for DNA Affinity Purification Assay (DAPA) ^56^. As for SNP-seq, 500 ng of different alleles (31 bp, biotinylated) from a SNPs were attached to 25 μl streptavidin-Dynabeads and then incubated with 100 μg of THP-1 nuclear extract for 1 h with or without non-biotinylated competitor (the same alleles). The incubation mixture was then washed PBST 5 times before adding elution buffer (2x Laemmli protein sample buffer, Bio-Rad, catalog number 1610747). The elution was then used for western blot.

### Western blot

Antibodies were diluted in PBST according to manufacturer’s suggestion. Antibodies used for western blot are anti-GFI1 antibody (Thermo Scientific, catalog number PA5-23495); anti-ATF3 antibody (Cell Signaling, catalog number 33593S), anti-FRA2 antibody (Abcam, catalog number ab124830), anti-TRAF1 antibody (Millipore Sigma, catalog number MABC260), anti-vinculin antibody (Bio-Rad, catalog number MCA465GA), goat anti-Rabbit IgG (H+L) cross-adsorbed secondary antibody, HRP (Thermo Scientific, catalog number G-21234) and goat anti-mouse IgG-HRP (Santa Cruz, catalog number sc-525408).

### CRISPR-cas9 knockout

FRA2 gene in THP-1 cells was knockout by CRISPR-cas9 technique through nucleofection (4D nucleofector from Lonza). 1 μl of 40 μM sgRNAs (two sgRNAs for each reaction to generate a deletion) for FRA2 gene were mixed with 1 μl of 20 μM spCas9 and incubated at 37 degrees for 15 minutes to form ribonucleoprotein (RNP). Formed RNP was kept on ice before use. THP-1 cells were then washed with PBS twice and resuspended in SG solution (Lonza, catalog number V4XC-3032), 200k cells per 20 μl solution. Next, RNP was added into each 20 μl solution with THP-1 cells and the whole mixture was transferred into nucleofection strip wells for nucleofection with program code FF100. After nucleofection, cells were transferred into pre-warmed culture medium and incubated for 72 hours before analyzing knockout efficiency. sgRNAs used are: 5’-GGAGAAGCGUCGCAUCCGGC-3’ and 5’-GAACCGACGCCGGGAGCUGA-3’ for k/o clone 1; 5’-CACCGCGGAUCAUGUACCAG-3’ and 5’-GCGCACGCCGAGUCCUACUC-3’ for k/o clone 2.

### RT-qPCR

Total RNA was isolated with Absolutely RNA Miniprep Kit (Agilent, catalog number 400800). cDNA was synthesized with AffinityScript QPCR cDNA Synthesis Kit (Agilent, catalog number 600559). All procedures were performed following the manufacturer’s protocols. RT-qPCR was done with a Agilent AriaMX qPCR machine according to the protocol for Brilliant II SYBR QPCR Low ROX Mstr Mx (Agilent, catalog number 600830). FRA2 qPCR primers: for k/o clone 1, forward, 5’-CAGCCAGCTTGTTCCTCT-3’, reverse, 5’-GATCAAGACCATTGGCACCA-3’; for k/o clone 2, forward, 5’-CAGCAGAAATTCCGGGTAGAT-3’, reverse, 5’-GGTATGGGTTGGACATGGAG-3’. TRAF1 qPCR primers: forward, 5’-GCCCTTCCGGAACAAGGTC-3’, reverse, 5’-CGTCAATGGCGTGCTCAC-3’.

### Chromatin Immunoprecipitation (ChIP)-qPCR

ChIP was performed for THP-1 cells and human monocytes using CHIP-it kit (Active motif, catalog number 53042) according to manufacturer’s protocols. Antibodies used are isotype control antibody and anti-FRA2 antibody from EMSA. The primers used for THP-1 cell CHIP-qPCR are forward, 5’-CTCCTCCTTTGTCATCATGTT-3’, reverse, 5’- TGGTCAGTTTCCTGGCAAATA-3’; primers and probes used for monocytes CHIP-qPCR are: forward 5’- AGCCTCTCCTCGCTATTC-3’, reverse 5’- GAAGGTGGCAAAGCTGAA-3’, A_Probe 5’- TG+A+C+GA+CAAAG+GA, B_Probe 5’- TG+A+T+GA+CAA+AG+GA.

### Allele discrimination CHIP-qPCR

CHIP was performed using human monocytes with rs7034653 A/G genotype. Antibodies used are isotype control antibody and anti-FRA2 antibody from EMSA. Probes and primers used are: G_Probe, 5’- TG+A+C+GA+CAAAG+GA, A_Probe, 5’- TG+A+T+GA+CAA+AG+GA; Forward primer, 5’- AGCCTCTCCTCGCTATTC, Reverse primer, 5’- GAAGGTGGCAAAGCTGAA.

### Conditional analysis on RA GWAS

We performed conditional analysis within ±1 Mb from the lead variant rs7034653 in each of the 25 EUR RA GWAS cohort, and meta-analyzed the 25 results using a fixed effect model, using methods and RA cohorts detailed in reference ^10^.

### Statistical analyses

All statistical analysis was done using GraphPad software (Prism) using one-way analysis of variance (ANOVA) for comparison of multiple groups, or unpaired t-test (non-parametric Mann-Whitney test) for two groups, with P-values as indicated in the figure legends.

## References

1. Farh, K.K., Marson, A., Zhu, J., Kleinewietfeld, M., Housley, W.J., Beik, S., Shoresh, N., Whitton, H., Ryan, R.J., Shishkin, A.A., et al. (2015). Genetic and epigenetic fine mapping of causal autoimmune disease variants. Nature 518, 337–343. 10.1038/nature13835.

2. Li, G., Martínez-Bonet, M., Wu, D., Yang, Y., Cui, J., Nguyen, H.N., Cunin, P., Levescot, A., Bai, M., Westra, H.J., et al. (2018). High-throughput identification of noncoding functional SNPs via type IIS enzyme restriction. Nat Genet 50, 1180–1188. 10.1038/s41588-018-0159-z.

3. Lu, X., Chen, X., Forney, C., Donmez, O., Miller, D., Parameswaran, S., Hong, T., Huang, Y., Pujato, M., Cazares, T., et al. (2021). Global discovery of lupus genetic risk variant allelic enhancer activity. Nature Communications 12, 1611. 10.1038/s41467-021-21854-5.

4. Moore, J.H., Asselbergs, F.W., and Williams, S.M. (2010). Bioinformatics challenges for genomewide association studies. Bioinformatics 26, 445–455. 10.1093/bioinformatics/btp713.

5. Zhang, Q., Bastard, P., Liu, Z., Le Pen, J., Moncada-Velez, M., Chen, J., Ogishi, M., Sabli, I.K.D., Hodeib, S., Korol, C., et al. (2020). Inborn errors of type I IFN immunity in patients with lifethreatening COVID-19. Science 370, eabd4570. doi:10.1126/science.abd4570.

6. Okada, Y., Wu, D., Trynka, G., Raj, T., Terao, C., Ikari, K., Kochi, Y., Ohmura, K., Suzuki, A., Yoshida, S., et al. (2014). Genetics of rheumatoid arthritis contributes to biology and drug discovery. Nature 506, 376–381. 10.1038/nature12873.

7. Stahl, E.A., Raychaudhuri, S., Remmers, E.F., Xie, G., Eyre, S., Thomson, B.P., Li, Y., Kurreeman, F.A.S., Zhernakova, A., Hinks, A., et al. (2010). Genome-wide association study meta-analysis identifies seven new rheumatoid arthritis risk loci. Nature Genetics 42, 508–514. 10.1038/ng.582.

8. Okada, Y., Terao, C., Ikari, K., Kochi, Y., Ohmura, K., Suzuki, A., Kawaguchi, T., Stahl, E.A., Kurreeman, F.A.S., Nishida, N., et al. (2012). Meta-analysis identifies nine new loci associated with rheumatoid arthritis in the Japanese population. Nature Genetics 44, 511–516. 10.1038/ng.2231.

9. Eyre, S., Bowes, J., Diogo, D., Lee, A., Barton, A., Martin, P., Zhernakova, A., Stahl, E., Viatte, S., McAllister, K., et al. (2012). High-density genetic mapping identifies new susceptibility loci for rheumatoid arthritis. Nature Genetics 44, 1336–1340. 10.1038/ng.2462.

10. Ishigaki, K., Sakaue, S., Terao, C., Luo, Y., Sonehara, K., Yamaguchi, K., Amariuta, T., Too, C.L., Laufer, V.A., Scott, I.C., et al. (2021). Trans-ancestry genome-wide association study identifies novel genetic mechanisms in rheumatoid arthritis. medRxiv, 2021.2012.2001.21267132. 10.1101/2021.12.01.21267132.

11. Plenge, R.M., Seielstad, M., Padyukov, L., Lee, A.T., Remmers, E.F., Ding, B., Liew, A., Khalili, H., Chandrasekaran, A., Davies, L.R.L., et al. (2007). TRAF1–C5 as a Risk Locus for Rheumatoid Arthritis — A Genomewide Study. New England Journal of Medicine 357, 1199–1209. 10.1056/NEJMoa073491.

12. Behrens, E.M., Finkel, T.H., Bradfield, J.P., Kim, C.E., Linton, L., Casalunovo, T., Frackelton, E.C., Santa, E., Otieno, F.G., Glessner, J.T., et al. (2008). Association of the TRAF1-C5 locus on chromosome 9 with juvenile idiopathic arthritis. Arthritis Rheum 58, 2206–2207. 10.1002/art.23603.

13. Albers, H.M., Kurreeman, F.A., Houwing-Duistermaat, J.J., Brinkman, D.M., Kamphuis, S.S., Girschick, H.J., Wouters, C., Van Rossum, M.A., Verduijn, W., Toes, R.E., et al. (2008). The TRAF1/C5 region is a risk factor for polyarthritis in juvenile idiopathic arthritis. Ann Rheum Dis 67, 1578–1580. 10.1136/ard.2008.089060.

14. Petty, R.E., Southwood, T.R., Manners, P., Baum, J., Glass, D.N., Goldenberg, J., He, X., Maldonado-Cocco, J., Orozco-Alcala, J., Prieur, A.-M., et al. (2004). International League of Associations for Rheumatology classification of juvenile idiopathic arthritis: second revision, Edmonton, 2001. The Journal of Rheumatology 31, 390–392.

15. McIntosh, L.A., Marion, M.C., Sudman, M., Comeau, M.E., Becker, M.L., Bohnsack, J.F., Fingerlin, T.E., Griffin, T.A., Haas, J.P., Lovell, D.J., et al. (2017). Genome-Wide Association Meta-Analysis Reveals Novel Juvenile Idiopathic Arthritis Susceptibility Loci. Arthritis Rheumatol 69, 2222–2232. 10.1002/art.40216.

16. Zapata, J.M., and Reed, J.C. (2002). TRAF1: lord without a RING. Sci STKE 2002, pe27. 10.1126/stke.2002.133.pe27.

17. Edilova, M.I., Abdul-Sater, A.A., and Watts, T.H. (2018). TRAF1 Signaling in Human Health and Disease. Frontiers in Immunology 9. 10.3389/fimmu.2018.02969.

18. Tsitsikov, E.N., Laouini, D., Dunn, I.F., Sannikova, T.Y., Davidson, L., Alt, F.W., and Geha, R.S. (2001). TRAF1 is a negative regulator of TNF signaling. enhanced TNF signaling in TRAF1-deficient mice. Immunity 15, 647–657. 10.1016/s1074-7613(01)00207-2.

19. Abdul-Sater, A.A., Edilova, M.I., Clouthier, D.L., Mbanwi, A., Kremmer, E., and Watts, T.H. (2017). The signaling adaptor TRAF1 negatively regulates Toll-like receptor signaling and this underlies its role in rheumatic disease. Nat Immunol 18, 26–35. 10.1038/ni.3618.

20. Hinks, A., Cobb, J., Marion, M.C., Prahalad, S., Sudman, M., Bowes, J., Martin, P., Comeau, M.E., Sajuthi, S., Andrews, R., et al. (2013). Dense genotyping of immune-related disease regions identifies 14 new susceptibility loci for juvenile idiopathic arthritis. Nat Genet 45, 664–669. 10.1038/ng.2614.

21. Fairfax, B.P., Humburg, P., Makino, S., Naranbhai, V., Wong, D., Lau, E., Jostins, L., Plant, K., Andrews, R., McGee, C., and Knight, J.C. (2014). Innate immune activity conditions the effect of regulatory variants upon monocyte gene expression. Science 343, 1246949. 10.1126/science.1246949.

22. Quach, H., Rotival, M., Pothlichet, J., Loh, Y.E., Dannemann, M., Zidane, N., Laval, G., Patin, E., Harmant, C., Lopez, M., et al. (2016). Genetic Adaptation and Neandertal Admixture Shaped the Immune System of Human Populations. Cell 167, 643–656 e617. 10.1016/j.cell.2016.09.024.

23. Alasoo, K., Rodrigues, J., Mukhopadhyay, S., Knights, A.J., Mann, A.L., Kundu, K., Hale, C., Dougan, G., Gaffney, D.J., and Consortium, H. (2018). Shared genetic effects on chromatin and gene expression indicate a role for enhancer priming in immune response. Nature Genetics 50, 424–431. 10.1038/s41588-018-0046-7.

24. Chen, L., Ge, B., Casale, F.P., Vasquez, L., Kwan, T., Garrido-Martin, D., Watt, S., Yan, Y., Kundu, K., Ecker, S., et al. (2016). Genetic Drivers of Epigenetic and Transcriptional Variation in Human Immune Cells. Cell 167, 1398–1414 e1324. 10.1016/j.cell.2016.10.026.

25. Weirauch, M.T., Cote, A., Norel, R., Annala, M., Zhao, Y., Riley, T.R., Saez-Rodriguez, J., Cokelaer, T., Vedenko, A., Talukder, S., et al. (2013). Evaluation of methods for modeling transcription factor sequence specificity. Nat Biotechnol 31, 126–134. 10.1038/nbt.2486.

26. Weirauch, M.T., Yang, A., Albu, M., Cote, A.G., Montenegro-Montero, A., Drewe, P., Najafabadi, H.S., Lambert, S.A., Mann, I., Cook, K., et al. (2014). Determination and inference of eukaryotic transcription factor sequence specificity. Cell 158, 1431–1443. 10.1016/j.cell.2014.08.009.

27. Nigrovic, P.A., Colbert, R.A., Holers, V.M., Ozen, S., Ruperto, N., Thompson, S.D., Wedderburn, L.R., Yeung, R.S.M., and Martini, A. (2021). Biological classification of childhood arthritis: roadmap to a molecular nomenclature. Nat Rev Rheumatol 17, 257–269. 10.1038/s41584-021-00590-6.

28. Foletta, V.C. (1996). Transcription factor AP-1, and the role of Fra-2. Immunol Cell Biol 74, 121–133. 10.1038/icb.1996.17.

29. Birnhuber, A., Crnkovic, S., Biasin, V., Marsh, L.M., Odler, B., Sahu-Osen, A., Stacher-Priehse, E., Brcic, L., Schneider, F., Cikes, N., et al. (2019). IL-1 receptor blockade skews inflammation towards Th2 in a mouse model of systemic sclerosis. European Respiratory Journal 54, 1900154. 10.1183/13993003.00154-2019.

30. Granet, C., and Miossec, P. (2004). Combination of the pro-inflammatory cytokines IL-1, TNF-alpha and IL-17 leads to enhanced expression and additional recruitment of AP-1 family members, Egr-1 and NF-kappaB in osteoblast-like cells. Cytokine 26, 169–177. 10.1016/j.cyto.2004.03.002.

31. Granet, C., Maslinski, W., and Miossec, P. (2004). Increased AP-1 and NF-kappaB activation and recruitment with the combination of the proinflammatory cytokines IL-1beta, tumor necrosis factor alpha and IL-17 in rheumatoid synoviocytes. Arthritis Res Ther 6, R190–198. 10.1186/ar1159.

32. McInnes, I.B., and Schett, G. (2007). Cytokines in the pathogenesis of rheumatoid arthritis. Nat Rev Immunol 7, 429–442. 10.1038/nri2094.

33. Radner, H., and Aletaha, D. (2015). Anti-TNF in rheumatoid arthritis: an overview. Wien Med Wochenschr 165, 3–9. 10.1007/s10354-015-0344-y.

34. Bradley, J.R. (2008). TNF-mediated inflammatory disease. J Pathol 214, 149–160. 10.1002/path.2287.

35. Gabay, C., Lamacchia, C., and Palmer, G. (2010). IL-1 pathways in inflammation and human diseases. Nat Rev Rheumatol 6, 232–241. 10.1038/nrrheum.2010.4.

36. Gutierrez-Arcelus, M., Ongen, H., Lappalainen, T., Montgomery, S.B., Buil, A., Yurovsky, A., Bryois, J., Padioleau, I., Romano, L., Planchon, A., et al. (2015). Tissue-specific effects of genetic and epigenetic variation on gene regulation and splicing. PLoS Genet 11, e1004958. 10.1371/journal.pgen.1004958.

37. Speiser, D.E., Lee, S.Y., Wong, B., Arron, J., Santana, A., Kong, Y.Y., Ohashi, P.S., and Choi, Y. (1997). A regulatory role for TRAF1 in antigen-induced apoptosis of T cells. J Exp Med 185, 1777–1783. 10.1084/jem.185.10.1777.

38. Xie, P., Hostager, B.S., Munroe, M.E., Moore, C.R., and Bishop, G.A. (2006). Cooperation between TNF receptor-associated factors 1 and 2 in CD40 signaling. J Immunol 176, 5388–5400. 10.4049/jimmunol.176.9.5388.

39. Wang, C.Y., Mayo, M.W., Korneluk, R.G., Goeddel, D.V., and Baldwin, A.S., Jr. (1998). NF-kappaB antiapoptosis: induction of TRAF1 and TRAF2 and c-IAP1 and c-IAP2 to suppress caspase-8 activation. Science 281, 1680–1683. 10.1126/science.281.5383.1680.

40. McPherson, A.J., Snell, L.M., Mak, T.W., and Watts, T.H. (2012). Opposing roles for TRAF1 in the alternative versus classical NF-κB pathway in T cells. J Biol Chem 287, 23010–23019. 10.1074/jbc.M112.350538.

41. Arron, J.R., Pewzner-Jung, Y., Walsh, M.C., Kobayashi, T., and Choi, Y. (2002). Regulation of the subcellular localization of tumor necrosis factor receptor-associated factor (TRAF)2 by TRAF1 reveals mechanisms of TRAF2 signaling. J Exp Med 196, 923–934. 10.1084/jem.20020774.

42. Hinks, A., Bowes, J., Cobb, J., Ainsworth, H.C., Marion, M.C., Comeau, M.E., Sudman, M., Han, B., Becker, M.L., Bohnsack, J.F., et al. (2017). Fine-mapping the MHC locus in juvenile idiopathic arthritis (JIA) reveals genetic heterogeneity corresponding to distinct adult inflammatory arthritic diseases. Ann Rheum Dis 76, 765–772. 10.1136/annrheumdis-2016-210025.

43. Chistiakov, D.A., Savost’anov, K.V., and Baranov, A.A. (2014). Genetic background of juvenile idiopathic arthritis. Autoimmunity 47, 351–360. 10.3109/08916934.2014.889119.

44. Prahalad, S., and Glass, D.N. (2008). A comprehensive review of the genetics of juvenile idiopathic arthritis. Pediatr Rheumatol Online J 6, 11. 10.1186/1546-0096-6-11.

45. Han, B., Diogo, D., Eyre, S., Kallberg, H., Zhernakova, A., Bowes, J., Padyukov, L., Okada, Y., González-Gay, M.A., Rantapää-Dahlqvist, S., et al. (2014). Fine mapping seronegative and seropositive rheumatoid arthritis to shared and distinct HLA alleles by adjusting for the effects of heterogeneity. Am J Hum Genet 94, 522–532. 10.1016/j.ajhg.2014.02.013.

46. Kurkó, J., Besenyei, T., Laki, J., Glant, T.T., Mikecz, K., and Szekanecz, Z. (2013). Genetics of rheumatoid arthritis - a comprehensive review. Clin Rev Allergy Immunol 45, 170–179. 10.1007/s12016-012-8346-7.

47. Terao, C., Brynedal, B., Chen, Z., Jiang, X., Westerlind, H., Hansson, M., Jakobsson, P.J., Lundberg, K., Skriner, K., Serre, G., et al. (2019). Distinct HLA Associations with Rheumatoid Arthritis Subsets Defined by Serological Subphenotype. Am J Hum Genet 105, 616–624. 10.1016/j.ajhg.2019.08.002.

48. Manichaikul, A., Mychaleckyj, J.C., Rich, S.S., Daly, K., Sale, M., and Chen, W.-M. (2010). Robust relationship inference in genome-wide association studies. Bioinformatics 26, 2867–2873. 10.1093/bioinformatics/btq559.

49. Alexander, D.H., Novembre, J., and Lange, K. (2009). Fast model-based estimation of ancestry in unrelated individuals. Genome Res 19, 1655–1664. 10.1101/gr.094052.109.

50. Langefeld, C.D., Ainsworth, H.C., Graham, D.S.C., Kelly, J.A., Comeau, M.E., Marion, M.C., Howard, T.D., Ramos, P.S., Croker, J.A., Morris, D.L., et al. (2017). Transancestral mapping and genetic load in systemic lupus erythematosus. Nature Communications 8, 16021. 10.1038/ncomms16021.

51. Kerimov, N., Hayhurst, J.D., Peikova, K., Manning, J.R., Walter, P., Kolberg, L., Samoviča, M., Sakthivel, M.P., Kuzmin, I., Trevanion, S.J., et al. (2021). A compendium of uniformly processed human gene expression and splicing quantitative trait loci. Nature Genetics 53, 1290–1299. 10.1038/s41588-021-00924-w.

52. Momozawa, Y., Dmitrieva, J., Théâtre, E., Deffontaine, V., Rahmouni, S., Charloteaux, B., Crins, F., Docampo, E., Elansary, M., Gori, A.-S., et al. (2018). IBD risk loci are enriched in multigenic regulatory modules encompassing putative causative genes. Nature Communications 9, 2427. 10.1038/s41467-018-04365-8.

53. Nédélec, Y., Sanz, J., Baharian, G., Szpiech, Z.A., Pacis, A., Dumaine, A., Grenier, J.C., Freiman, A., Sams, A.J., Hebert, S., et al. (2016). Genetic Ancestry and Natural Selection Drive Population Differences in Immune Responses to Pathogens. Cell 167, 657–669.e621. 10.1016/j.cell.2016.09.025.

54. Schmiedel, B.J., Singh, D., Madrigal, A., Valdovino-Gonzalez, A.G., White, B.M., Zapardiel-Gonzalo, J., Ha, B., Altay, G., Greenbaum, J.A., McVicker, G., et al. (2018). Impact of Genetic Polymorphisms on Human Immune Cell Gene Expression. Cell 175, 1701–1715.e1716. 10.1016/j.cell.2018.10.022.

55. Giambartolomei, C., Vukcevic, D., Schadt, E.E., Franke, L., Hingorani, A.D., Wallace, C., and Plagnol, V. (2014). Bayesian test for colocalisation between pairs of genetic association studies using summary statistics. PLoS Genet 10, e1004383. 10.1371/journal.pgen.1004383.

56. Miller, D.E., Patel, Z.H., Lu, X., Lynch, A.T., Weirauch, M.T., and Kottyan, L.C. (2016). Screening for Functional Non-coding Genetic Variants Using Electrophoretic Mobility Shift Assay (EMSA) and DNA-affinity Precipitation Assay (DAPA). J Vis Exp. 10.3791/54093.

