## supplemental figure 1 for "Identification of a regulatory pathway governing TRAF1 via an arthritis-associated non-coding variant"

### S. Figure 1

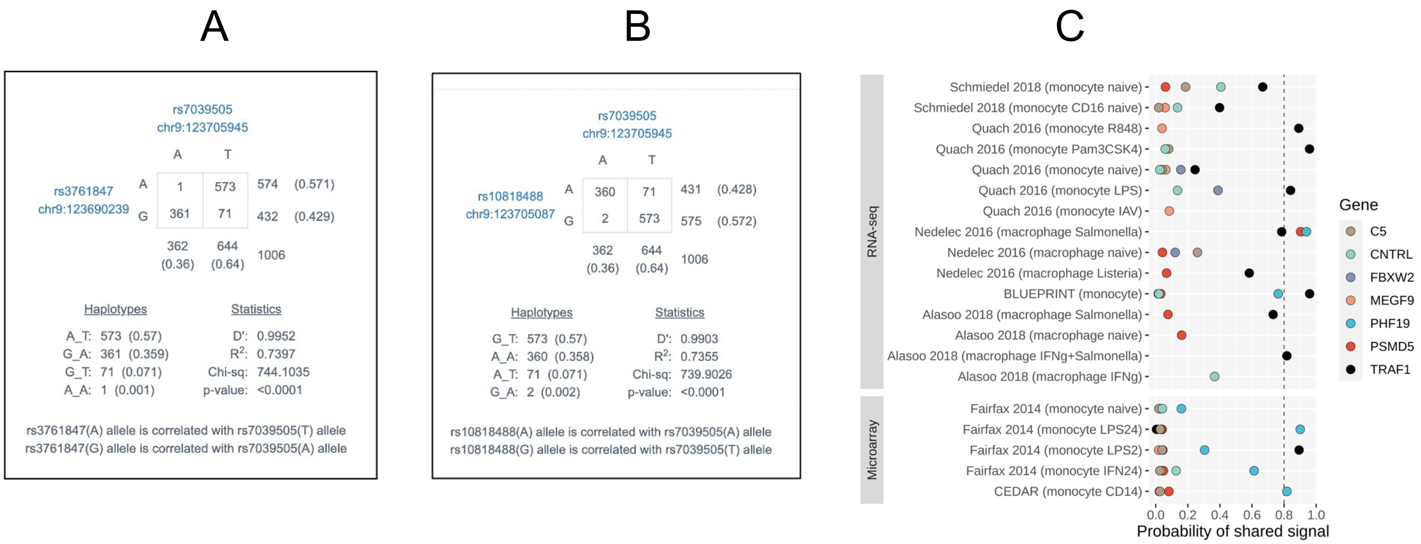


**S. Figure 1.** LD pair of rs7039505 with other variants associated with risk of RA (rs3761847, **(A)**) and JIA (rs10818488, **(B)** ) identified previously. **(C)** Colocalization analysis for the JIA ImmunoChip variants in the TRAF1-C5 locus against the eQTL Catalogue in the monocytes studies. TRAF1 is the major gene with a posterior probability (PP4) > 0.8.
