## supplemental figure 2 for "Identification of a regulatory pathway governing TRAF1 via an arthritis-associated non-coding variant"

### S. Figure 2

A

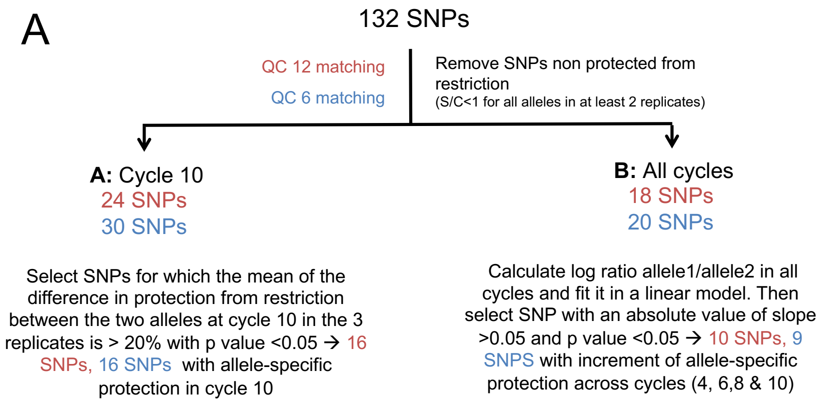


B

| QC 12 matching | | QC 6 matching | | Score |
|---|---|---|---|---|
| Aproach A | Aproach B | Aproach A | Aproach B | |
| rs7021880 | rs7021880 | rs7021880 | rs7021880 | **4** |
| rs7034653 | rs7034653 | rs7034653 | rs7034653 | **4** |
| rs758959 | rs758959 | rs758959 | rs758959 | **4** |
| rs6478484 | rs6478484 | rs6478484 | rs6478484 | **4** |
| rs7858209 | | rs7858209 | rs7858209 | **3** |
| rs3761849 | | rs3761849 | rs3761849 | **3** |
| rs7875829 | rs7875829 | | | **2** |
| rs9886724 | rs9886724 | | | **2** |
| rs10760129 | rs10760129 | | | **2** |
| | | rs10985073 | rs10985073 | **2** |
| | | rs1609810 | rs1609810 | **2** |
| rs1008381 | | rs1008381 | | 2 |
| rs10435844 | | rs10435844 | | 2 |
| rs7021206 | | rs7021206 | | 2 |
| | rs10760130 | | rs10760130 | 2 |
| | | rs10117059 | | 1 |
| | | rs10739579 | | 1 |
| | | rs1468671 | | 1 |
| | | rs1860823 | | 1 |
| | | rs4837804 | | 1 |
| rs10985102 | | | | 1 |
| rs7037195 | | | | 1 |
| rs2109896 | | | | 1 |
| rs4837799 | | | | 1 |
| | rs10739577 | | | 1 |
| | rs10739581 | | | 1 |

**S. Figure 2. Selection of SNPs for experimental investigation from SNP-seq. (A)** Detailed data analysis approaches for SNP-seq NGS data. QC6 and QC12 matching means 6 or 12 nucleotides on each side of SNP site from the NGS sequencing data matched to the original sequence. **(B)** Summary table of all the SNPs selected from SNP-seq based on one of the analysis approaches. The 11 SNPs highlighted in yellow were chosen for downstream validation.
