## supplemental figure 3 for "Identification of a regulatory pathway governing TRAF1 via an arthritis-associated non-coding variant"

### S. Figure 3

A

#### rs7034653 cis-eQTLs with TRAF1 in monocyte

Monocyte

Raw P value:0.0098 Adj. P value: 0.9 Effect size: 0.4

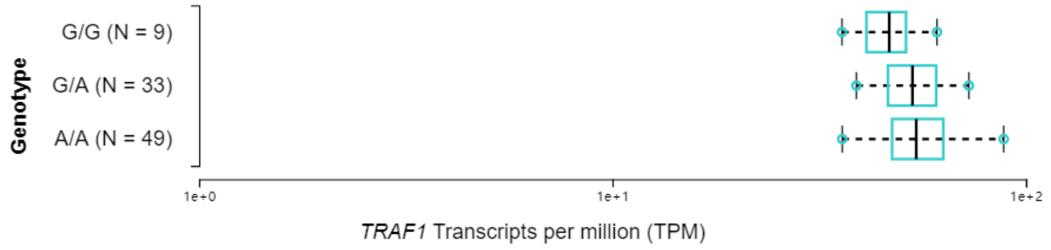


#### rs1609810 cis-eQTLs with TRAF1 in monocyte

Monocyte

Raw P value:0.008 Adj. P value: 0.86 Effect size: 0.41

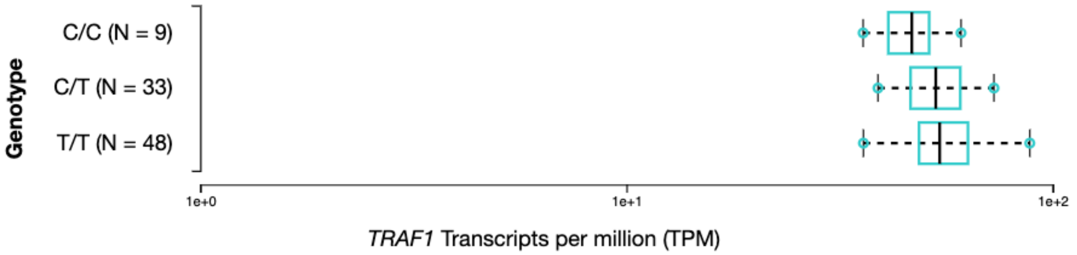


B

#### SNPs in cis-mQTLs with TRAF1 from BIOS QTL browser

| P-value | SNP | SNP Chr. | SNP Chr. Position | CpG | CpG Chr. | CpG Chr. position | SNP Alleles | Assesed Allele | Z-score | Gene name | FDR |
|---|---|---|---|---|---|---|---|---|---|---|---|
| 6.15E-229 | rs7034653 | 9 | 123687372 | cg21152671 | 9 | 123690610 | A/G | G | 32.3 | TRAF1 | 0 |
| 5.55E-76 | rs1014530 | 9 | 123685092 | cg14064762 | 9 | 123688769 | C/T | T | 18.45 | TRAF1 | 0 |
| 6.03E-56 | rs1014530 | 9 | 123685092 | cg15551881 | 9 | 123688691 | C/T | T | 15.76 | TRAF1 | 0 |
| 1.66E-13 | rs77617771 | 9 | 123729896 | cg15551881 | 9 | 123688691 | G/A | A | 7.37 | TRAF1 | 0 |
| 2.49E-06 | rs10985143 | 9 | 123829010 | cg15551881 | 9 | 123688691 | C/G | G | 4.71 | TRAF1 | 0 |
| 3.14E-05 | rs10739574 | 9 | 123594836 | cg14064762 | 9 | 123688769 | G/A | A | 4.16 | TRAF1 | 0.02 |

**S. Figure 3. (A)** Expression quantitative trait loci data for rs7034653 and rs1609810 in human monocytes. **(B)** SNPs in cis-methylation quantitative trait loci with TRAF1 from BIOS QTL browser.
