## supplemental figure 4 for "Identification of a regulatory pathway governing TRAF1 via an arthritis-associated non-coding variant"

### S. Figure 4

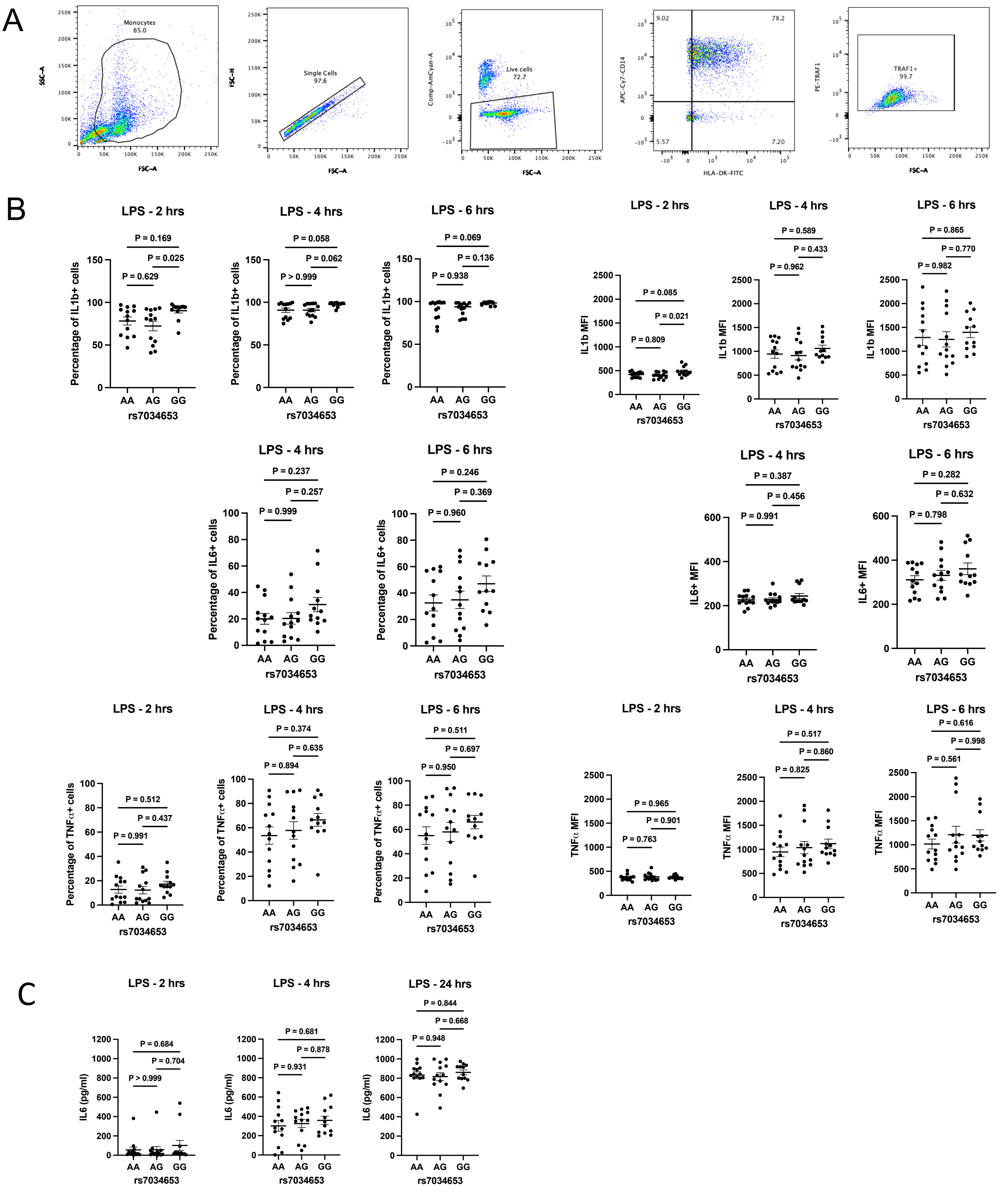


**S. Figure 4. Cytokine production in human monocytes as a function of genotype at rs7034653. (A)** Gating strategy for human monocytes. **(B)** Purified monocytes from healthy human PBMCs were treated with LPS (100 ng/ml for 2, 4 or 6 hrs), and percentage of cytokine producing cells as well as MFI was assessed by intracellular staining for IL-1, IL-6, and TNF. (C) IL-6 secreted by LPS-treated human monocytes measured by ELISA. Each symbol represents one donor; n = 13, 13, 12 human donors of each genotype AA, AG, GG, respectively. Statistical analysis was performed using one-way ANOVA multiple comparisons, all p values are shown in the figures.
